# Parallel holographic illumination enables sub-millisecond two-photon optogenetic activation in mouse visual cortex *in vivo*

**DOI:** 10.1101/250795

**Authors:** I-Wen Chen, Emiliano Ronzitti, Brian R. Lee, Tanya L. Daigle, Hongkui Zeng, Eirini Papagiakoumou, Valentina Emiliani

## Abstract

Selective control of action potential generation in individual cells from a neuronal ensemble is desirable for dissecting circuit mechanisms underlying perception and behavior. Here, by using two-photon (2P) temporally focused computer-generated holography (TF-CGH), we demonstrate optical manipulation of neuronal excitability at the supragranular layers of anesthetized mouse visual cortex. Utilizing amplified laser-pulses delivered via a localized holographic spot, our optical system achieves suprathreshold activation by exciting either of the three optogenetic actuators, ReaChR, CoChR or ChrimsonR, with brief illumination (≤ 10 ms) at moderate excitation power ((in average ≤ 0.2 mW/µm^2^ corresponding to ≤ 25 mW/cell). Using 2P-guided whole-cell or cell-attached recordings in positive neurons expressing respective opsin *in vivo*, we find that parallel illumination induces spikes of millisecond temporal resolution and sub-millisecond precision, which are preserved upon repetitive illuminations up to tens of Hz. Holographic stimulation thus enables temporally precise optogenetic activation independently of opsin’s channel kinetics. Furthermore, we demonstrate that parallel optogenetic activation can be combined with functional imaging for all-optical control of a neuronal sub-population that co-expresses the photosensitive opsin ReaChR and the calcium indicator GCaMP6s. Parallel optical control of neuronal activity with cellular resolution and millisecond temporal precision should be advantageous for investigating neuronal connections and further yielding causal links between connectivity, microcircuit dynamics, and brain functions.

**Significance statement:** Recent development of optogenetics allows probing the neuronal microcircuit with light by optically actuating genetically-encoded light-sensitive opsins expressed in the target cells. Here, we apply holographic light shaping and temporal focusing to simultaneously deliver axially-confined holographic patterns to opsin-positive cells situated in the living mouse cortex. Parallel illumination efficiently induces action potentials with high temporal resolution and precision for three opsins of different kinetics. We demonstrated all-optical experiments by extending the parallel optogenetic activation at low intensity to multiple neurons and concurrently monitoring their calcium dynamics. These results demonstrate fast and temporally precise *in vivo* control of a neuronal sub-population, opening new opportunities to reveal circuit mechanisms underlying brain functions.

## Introduction

The coordinated spike timing between neurons at millisecond precision underlies various synaptic mechanisms which could play significant roles in regulating sensation, perception, and cognitive function (1, 2). Optogenetics with its still-expanding genetic toolbox (3–6), opened the way to investigate those mechanisms with all-optical approaches (7–12). However, reaching the necessary precision to replicate and monitor neuronal circuits’ activity with light *in viv*o is still challenged by the requirement for photostimulating one or several individually chosen cells within scattering tissues with millisecond precision while concurrently monitoring the evoked activity.

Wide-field illumination using visible light has proved the capability of driving neuronal activity with such temporal precision reaching high spiking rate, with millisecond peak latencies and sub-millisecond jitter (i.e. the standard deviation of latencies) in cultured cells and acute brain slices. For example, for ChR2 and recently developed opsins ReaChR, Chronos and ChrimsonR, trains of action potentials (AP) up to tens of Hz can be induced following repetitive illumination of brief one-photon (1P) light-pulses ≤ 5 ms, and single spike of peak latency < 10 ms and jitter < 1 ms can be generated (5, 6). However, 1P illumination lacks optical sectioning and, in scattering samples, penetration depth. As a consequence, its use for *in vivo* activity manipulation with cellular resolution has been limited so far to shallow depths or transparent animals (13–15).

Two-photon (2P) excitation provides accurate targeting of neurons and optical sectioning, as well as the prerequisite of reduced scattering in tissue (16, 17). In particular, 2P optogenetic neuronal activation has been realized by employing either, or a hybrid of two approaches: the scanning and the parallel method. The former involves deflecting a focused beam across the target soma (18–20), whereas the later engages a light-pattern covering simultaneously the entire cell soma. Parallel methods for 2P optogenetic activation so far employ phase-modulation methods such as computer-generated holography (CGH) (21–23), generalized phase contrast (GPC) (24–26), the use of large Gaussian beams, extended to fit the size of a soma by decreasing the effective numerical aperture (NA) of the microscope objective (27, 28), or by defocusing the original beam (29). Axial confinement of light-patterns can be achieved and preserved till hundreds of µm by integrating these approaches with temporal focusing (TF) (22, 25–31).

Until now, *in vivo* 2P optogenetic activation in the mouse brain has been only reported for the opsin C1V1 by using scanning-based excitation, demonstrating AP generation of latencies 20-35 ms and jitter 6-20 ms (20). Spike induction by using temporally focused Gaussian beams has been shown for opsins C1V1 (27) and ChrimsonR (29), but its achievable temporal resolution and precision have not yet been specified.

Here, by combining TF with CGH, we demonstrated *in vivo* 2P activation in single or multiple neurons expressing either of the three opsins, ReaChR, CoChR, and ChrimsonR (5, 6) and characterizing their achievable temporal resolution and precision. The more red-shifted excitation spectral peaks of ReaChR and ChrimsonR render them attractive candidates for dual-channel photostimulation or all-optical manipulation by co-expressing either another blue-shifted opsin or an activity probe of green-shifted excitation spectra such as GCaMP (32). On the other hand, 2P parallel illumination efficiently activates the opsin CoChR, as well as its soma-targeting variant (33), thus holding promises to be combined with red-shifted activity sensor such as RCaMP (34).

The three opsins differ in their photocycle properties of channel opening, closing and inactivating (7, 35), which may play important roles in the temporal properties of 2P activation. ReaChR is a slow opsin which displays the longest transition time spans between photocycle states, CoChR the intermediate, and ChrimsonR the fast (their respective off-time constants being ~90 ms, ~30 ms and ~15 ms) (6, 33, 36, 37).

Using 2P-guided whole-cell or cell-attached recordings to measure spikes induced by brief patterned illumination (≤ 10 ms) from pyramidal cells and interneurons in the anesthetized mouse V1, we estimated for the three opsins the temporal resolution and precision based on the peak latencies of induced APs. We found the condition of 2P parallel illumination which enables reliable and temporally precise suprathreshold activation with <1 ms jitter, for the 3 opsins of different channel kinetics. Finally, by co-expressing the green calcium sensor GCaMP6 with ReaChR, we demonstrated simultaneous multi-cell activation in an all-optical setting.

## Results

### Parallel 2P activation of neurons expressing ReaChR, CoChR, and ChrimsonR *in vivo*

To achieve scan-less optogenetic activation in the living mouse brain, we integrated CGH and TF with a 2P scanning microscope (Fig. S1A). Laser light-pulses at a repetition rate of 500 kHz were delivered through a 12-µm-diameter holographic spot (Fig. S1B) placed over the target cell soma. We investigated the 2P stimulation parameters necessary for inducing an AP in V1 cells 5-12 weeks after viral infection of the three opsins ReaChR, CoChR and ChrimsonR (see Materials and Methods). The power density of the excitation laser was increased until the target cell elicited an AP, which was recorded via 2P-guided whole-cell or cell-attached recordings (Fig. 1A). The temporal resolution and precision of AP induction were estimated respectively as the peak latencies’ arithmetic mean and standard deviation (i.e. the jitter). According to the spiking patterns of positive cells situated at L2/3 (117±3 µm from the brain surface; mean±s.e.m.), we identified that recordings for ReaChR (15/15 cells), CoChR (13/13 cells) and the majority of ChrimsonR (20/24 cells) were from putative excitatory pyramidal neurons, whereas few ChrimsonR recordings (4/24 cells) from putative fast-spiking interneurons. Using brief pulse illumination of 2, 3, 5, and 10 ms, we found the threshold power density in the range of 0.1-0.4 mW/µm^2^ (measured at the tip of objective, corresponding to 11-45 mW/cell), sufficient for triggering an AP with average latency less than 10 ms and jitter smaller than 2 ms for the 3 opsins (Fig. 1A and Table 1). By slightly increasing the excitation power above the threshold level, we obtained shortened AP latencies (< 9 ms) as well as sub-millisecond jitter (Fig. 1B-C, Fig. S2, and Table 2).

**Figure 1.**
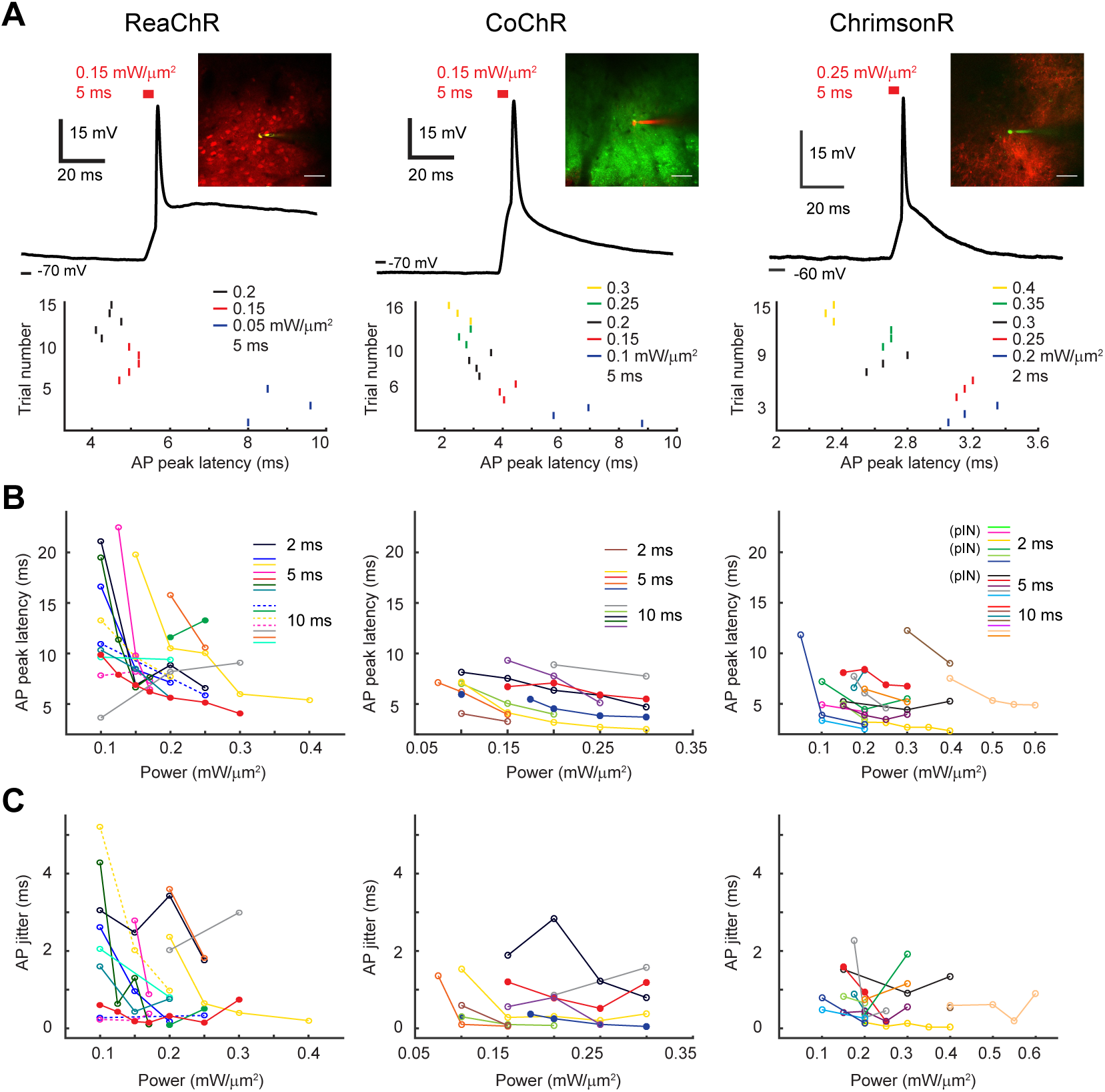
Precise 2P holographic activation of single cell *in vivo*. (A) Top: Example traces of AP induction at the L2/3 of anesthetized mouse visual cortex upon brief pulse of holographic illumination in cells expressing ReaChR, CoChR, and ChrimsonR respectively. Insets are 2P images of a target cell recorded via a glass pipette filled with Alexa Fluor 488 (green) or Alexa Fluor 592 (red). Bottom: Example raster plots of spike timing for each cell in response to different excitation power densities. Scale bar 47 μm. (B) AP peak latency in individual cells (connected dots represent data from the same cell) in relation to the excitation power density upon illumination of 2, 5, or 10 ms (n=10, 9, 11 for ReaChR, CoChR, and ChrimsonR). Solid circles indicate whole-cell recordings and open circles cell-attached recordings. For ChimsonR, 3 recordings were obtained from putative fast-spiking interneurons (pIN). (C) Jitter of AP peak latencies as a function of illumination power densities from the same cells as in (B).

**Table 1.** Spiking properties upon 2P pulse stimulation with threshold power density.

| Opsin | Light - pulse duration (ms) | AP latency (ms) | AP jitter (ms) | AP count | AP probability (%) | Laser power density** (mW/$\mu\text{m}^2$) | Estimated laser power density*** (mW/$\mu\text{m}^2$) |
| --- | --- | --- | --- | --- | --- | --- | --- |
| ReaChR | 10 | 9.48±1.41 (n=8) | 1.84±0.73 (n=7) | 0.98±0.065 (n=8) | 90.58±6.30 (n=8) | 0.14±0.018 (n=8) | 0.072±0.0083 (n=8) |
| | 5 | 11.16±2.42 (n=6) | 1.82±0.60 (n=6) | 1.11±0.070 (n=6) | 100.00±0.00 (n=6) | 0.13±0.017 (n=6) | 0.064±0.0065 (n=6) |
|  | 2 | 21.10 (n=1) | 3.05 (n=1) | 1.00 (n=1) | 100.00 (n=1) | 0.10 (n=1) | 0.054 (n=1) |
| CoChR | 10 | 8.18±0.58 (n=7) | 0.58±0.13 (n=6) | 1.00±0.072 (n=7) | 95.24±4.76 (n=7) | 0.19±0.038 (n=7) | 0.10±0.018 (n=7) |
| | 5 | 6.74±0.28 (n=4) | 1.10±0.27 (n=4) | 1.08±0.083 (n=4) | 100.00±0.00 (n=4) | 0.11±0.016 (n=4) | 0.056±0.0090 (n=4) |
|  | 3 | 6.93 (n=1) | 0.36 (n=1) | 1.00 (n=1) | 100.00 (n=1) | 0.15 (n=1) | 0.058 (n=1) |
|  | 2 | 4.05 (n=1) | 0.60 (n=1) | 1.00 (n=1) | 100.00 (n=1) | 0.10 (n=1) | 0.049 (n=1) |
| ChrimsonR | 10 | 8.50±0.74 (n=10) | 0.67±0.23 (n=8) | 0.89±0.054 (n=10) | 89.33±5.46 (n=10) | 0.24±0.021 (n=10) | 0.13±0.015 (n=10) |
| | 5 | 6.21±0.72 (n=6) | 1.30±0.26 (n=6) | 0.99±0.082 (n=6) | 93.33±5.77 (n=6) | 0.15±0.012 (n=6) | 0.080±0.0066 (n=6) |
| | 2 | 5.24±1.06 (n=8) | 0.30±0.13 (n=5) | 0.96±0.12 (n=8) | 87.5±6.10 (n=8) | 0.15±0.021 (n=8) | 0.073±0.011 (n=8) |
| ChrimsonR Tg | 15 | 13.32±0.75 (n=2) | 3.16±1.85 (n=2) | 0.82±0.017 (n=2) | 81.67±1.67 (n=2) | 0.38±0.075 (n=2) | 0.15±0.026 (n=2) |
| | 10 | 9.55±0.66 (n=5) | 2.39±1.08 (n=4) | 0.82±0.11 (n=5) | 77.00±10.20 (n=5) | 0.22±0.012 (n=5) | 0.11±0.013 (n=5) |
|  | 5 | 5.70 (n=1) | 1.15 (n=1) | 0.75 (n=1) | 75.00 (n=1) | 0.15 (n=1) | 0.070 (n=1) |

\* mean ± s.e.m.
\*\* Laser power density measured at the tip of objective
\*\*\* Estimated power density by assuming the scattering length at 1030 nm as 175 $\mu\text{m}$ , which is derived from simulations based on experimental data (26).

**Table 2.** Spiking properties upon 2P pulse stimulation with sub-millisecond precision.

| Opsin | Light-pulse duration (ms) | AP latency (ms) | AP jitter (ms) | AP count | AP probability (%) | Laser power density** (mW/$\mu\text{m}^2$) | Estimated laser power density*** (mW/$\mu\text{m}^2$) |
| --- | --- | --- | --- | --- | --- | --- | --- |
| ReaChR | 10 (n=7) | 8.80±0.77 | 0.58±0.15 | 1.24±0.19 | 100.00±0.00 | 0.16±0.020 | 0.081±0.0092 |
| | 5 (n=4) | 8.31±1.68 | 0.47±0.098 | 1.50±0.29 | 100.00±0.00 | 0.14±0.021 | 0.073±0.0087 |
| CoChR | 10 (n=7) | 7.69±0.77 | 0.61±0.12 | 1.05±0.047 | 100.00±0.00 | 0.22±0.038 | 0.12±0.018 |
| | 5 (n=4) | 5.84±0.62 | 0.37±0.15 | 0.92±0.083 | 91.67±8.33 | 0.14±0.024 | 0.072±0.014 |
|  | 3 (n=1) | 6.93 | 0.36 | 1.00 | 100.00 | 0.15 | 0.058 |
|  | 2 (n=1) | 4.05 | 0.60 | 1.00 | 100.00 | 0.10 | 0.049 |
| ChrimsonR | 10 (n=8) | 7.63±0.55 | 0.48±0.10 | 0.95±0.05 | 95.00±5.00 | 0.25±0.033 | 0.13±0.023 |
| | 5 (n=5) | 5.41±0.87 | 0.60±0.13 | 0.95±0.05 | 95.00±5.00 | 0.19±0.033 | 0.098±0.015 |
| | 2 (n=8) | 3.82±0.24 | 0.35±0.10 | 1.11±0.084 | 100.00±0.00 | 0.18±0.013 | 0.088±0.0072 |
| ChrimsonR Tg | 15 (n=1) | 13.45 | 0.63 | 1.00 | 100.00 | 0.40 | 0.16 |
| | 10 (n=4) | 8.57±0.77 | 0.66±0.19 | 1.00±0.00 | 100.00±0.00 | 0.29±0.038 | 0.15±0.020 |
|  | 5 (n=1) | 4.58 | 0.10 | 1.00 | 100.00 | 0.20 | 0.094 |

Further, using the transgenic mice of Cux2-CreERT2;Ai167 (38–40), in which ChrimsonR-tdTomato was specifically expressed on the cell membrane of L2/3 neurons, APs were induced by shining short light-pulses of 5, 10, or 15 ms with excitation intensity in the range of 0.15-0.45 mW/µm^2^, i.e., 17-51 mW/cell (138±15 µm deep, n=8; Fig. S3A). As before, we observed a shortening of AP latency and jitter (till < 1 ms) with the increasing excitation intensity. Compared to viral delivery of ChrimsonR, higher excitation power density and longer illumination duration were required for reaching AP threshold (p=0.065 and p=0.0034, 1-way ANOVA for preparation type) and obtaining sub-millisecond jitter (p=0.035 and p=0.015, 1-way ANOVA). The higher excitation power and longer illumination duration for precise suprathreshold activation may be resulted from the overall lower expression level of ChrimsonR in transgenic mice.

These results suggest that parallel optogenetic excitation, utilizing a low-repetition rate laser, achieves efficient membrane integration using low average power, thus enabling *in vivo* AP generation with < 1 ms jitter independently of the opsin channel kinetics, both for viral and transgenic expression of opsins.

### Temporally precise photostimulation of a train of patterned light-pulses

We investigated the stimulation conditions for precisely inducing a train of APs. Using illumination conditions according to the threshold power density of each cell, we examined the spiking properties upon a train of illuminations at different frequencies *in vivo* (Fig. 2A). For neurons expressing ReaChR, CoChR, and ChrimsonR via both viral and transgenic delivery upon illumination of 5 consecutive pulses, their average firing rates (FR) rose with the increasing stimulation frequencies of 10, 20 and 40 Hz (Fig. S3B and S4). Of note, membrane potential remained the most depolarized between light-pulses for ReaChR, the intermediate for CoChR, and the least for ChrimsonR (membrane potential after filtering out APs: 20.7±7.9, 18.7±3.3, 4.1±2.2 mV for ReaChR, CoChR, ChrimsonR upon 20-Hz photostimulation, n=3, 4, 2). The different degrees of repolarization between light-pulses are related to the different off-kinetics of the 3 opsins, with ChrimsonR the fastest, CoChR the middle, and ReaChR the slowest (6, 33, 36). For ChrimsonR, the fast repolarization between successive pulses enabled generation of AP train at even higher frequencies (> 40 Hz, n=2; Fig. S3B).

**Figure 2.**
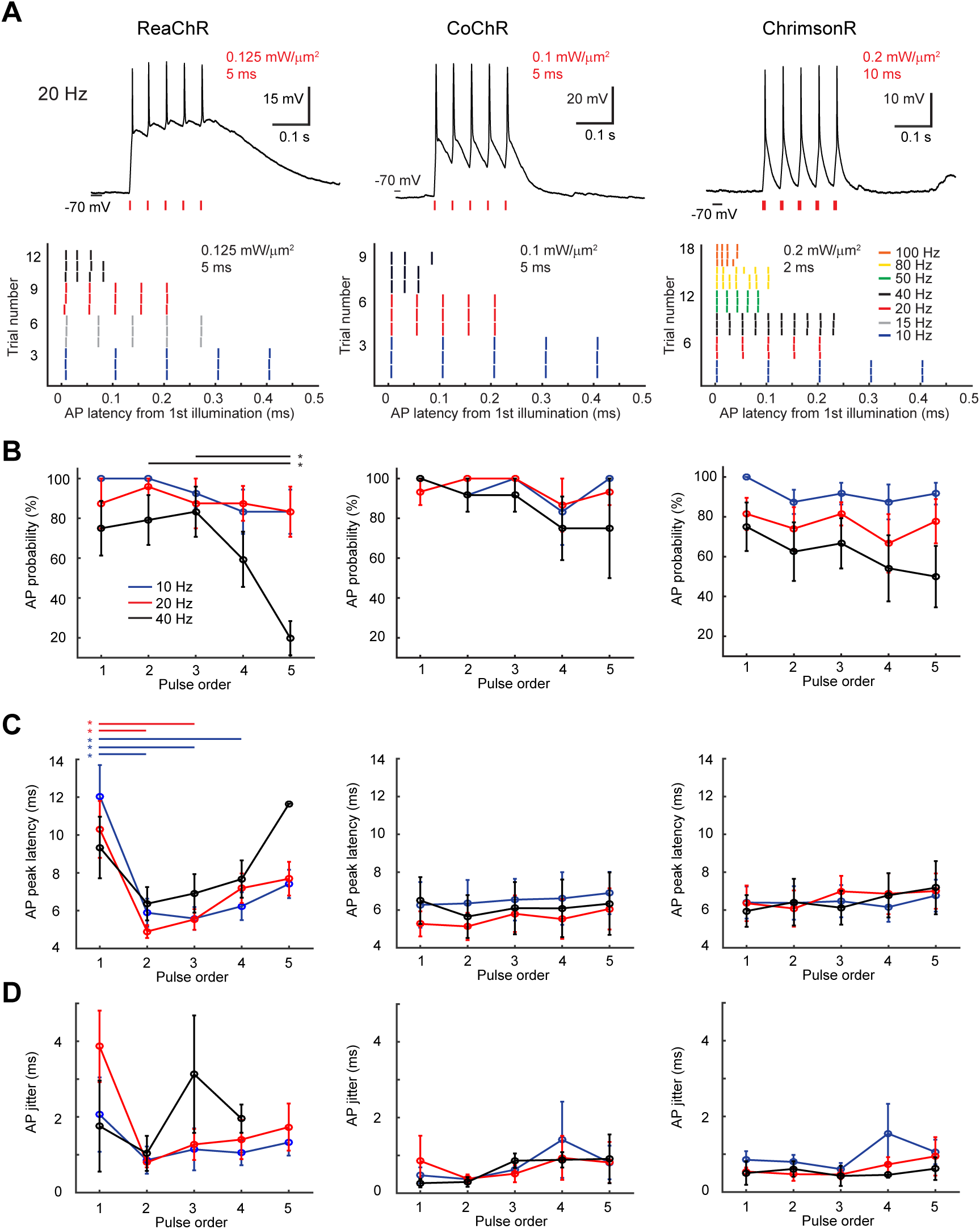
Precise generation of multiple APs with a train of patterned light-pulses. (A) Top: Representative whole-cell recordings of suprathreshold activation in vivo upon photo-stimulation of 5 illuminations at 20 Hz in ReaChR-, CoChR-, or ChrimsonR-positive neurons. Excitation intensity and light-pulse duration are indicated in red. Bottom: Example raster plots of AP peak latencies relative to the onset of first light-pulse from 5 or 10 illuminations at different frequencies. Photostimulation condition indicated in black. (B-D) AP probability, AP latency and jitter in relation to the pulse order of 5 illuminations at 10, 20, and 40 Hz for ReaChR, CoChR, and ChrimsonR respectively (mean±s.e.m.; n=9, 8, 8 for ReaChR in response to 10, 20, 40 Hz photostimulation, n=4, 5, 4 for CoChR, and n=8, 9, 8 for ChrimsonR). Note the significant decrease in AP latency and jitter upon the second stimulation for ReaChR. Asterisks denote significant difference for multiple comparison between pulse order.

For the 3 opsins, the probability of generating an AP decreased with the repeating occurrence of light-pulses at the 3 stimulation frequencies. This effect is more pronounced for ReaChR of slow channel kinetics. Upon photostimulation at a high frequency of 40 Hz, the AP probability in response to the 5-th illumination was significantly reduced for ReaChR (19.8±8.6%, n=8) whereas that remained ≥ 50% for CoChR (75.0±25.0%, n=4) and ChrimsonR (50.0±15.4%, n=8) (Fig. 2B).

The opsin photocycle also affected light-induced AP properties of peak latency and jitter upon repetitive illumination at 10, 20 and 40 Hz (p=0.0012 and p<0.0001, 3-way ANOVA for opsin type, pulse order and stimulation frequency) (Fig. 2C-D). Compared to the first illumination, the second light-pulse induced significantly shortened AP latencies in ReaChR-positive cells (p=0.0039 for pulse order in 3-way ANOVA, p<0.05 for 10 and 20 Hz in multiple comparisons). The slow off-kinetics of ReaChR may keep the cell membrane potential depolarized after the first light-induced AP for a longer period; in such an excited state, the second illumination may induce AP with increased temporal resolution of shortened peak latency.

To test the capability of our system to replicate, with < 1 ms precision, physiological activity patterns, we stimulated ChrimsonR-expressing cells in transgenic mouse with an illumination sequence reproducing the same temporal pattern of the target cell’s own spontaneous firing. Photoevoked spikes were played back following the original time course with sub-millisecond jitter (Fig. S5).

In sum, we demonstrated that following multiple parallel holographic illuminations, positive cells elicited a train of spikes of up to 40 Hz, with a < 10 ms latency and < 2 ms jitter, the overall temporal resolution and precision depending, as expected, on the opsin photocycle properties. This temporal precision enabled precise playback of neuronal spontaneous activity.

### Spatially precise photoactivation

Because of the clear visualization of cytosol fluorophore in ReaChR-expressing cells (Fig. 1A), we chose to map the photoactivation spatial selectivity of holographic activation *in vivo* in mouse V1 infected with the ReaChR-dTomato construct. To characterize the photoactivation axial selectivity, we recorded the spiking properties of target cell upon pulse stimulation at the threshold power density for each axial position after mechanically moving the objective out of focal plane. The estimated AP probabilities were obtained by applying two exponential fits on the average AP probabilities above and below the focal plane (Fig. 3B). Assuming the target soma’s shape is of a 12-µm-diameter sphere (grey-shaded area in Fig. 3B), the target cell fired AP of ~56% probability when the holographic spot was placed adjacent to the cell (axial positions of spot center ±12 µm) (Fig. 3A-B). The evoked AP probability further decreased when the excitation pattern was moved outside the target cell (axial positions >12 µm or <-12 µm). Photoactivation axial selectivity was 39 µm as the full-width at half-maximum (FWHM) computed from the estimated AP probabilities along the z-axis. The broad photoactivation axial selectivity measured *in vivo* cannot be explained by convolving the optical axial resolution with the cell size (Fig. S6A; see Materials and Methods), and may be resulted from activation of neurites by the defocused light (33, 30). Tissue scattering may contribute to the larger errors in AP probabilities deeper below the focal plane. In addition, light-induced APs in some cells (4/12) displayed lengthened latencies and enlarged jitter when the objective moved axially away from the focal plane (Fig. S6B). Upon defocused illumination, the temporal properties of AP generation in the target cell may be modulated by backpropagation of its dendritic activation or integration of dendritic signaling from neighboring neurons (41). Finally, we evaluated the lateral photoactivation selectivity by measuring the spiking properties after placing the holographic spot laterally away from the target soma. When the center of excitation volume was placed at a lateral position of 7 µm, the target neuron fired AP of ~50% (Fig. 3C).

**Figure 3.**
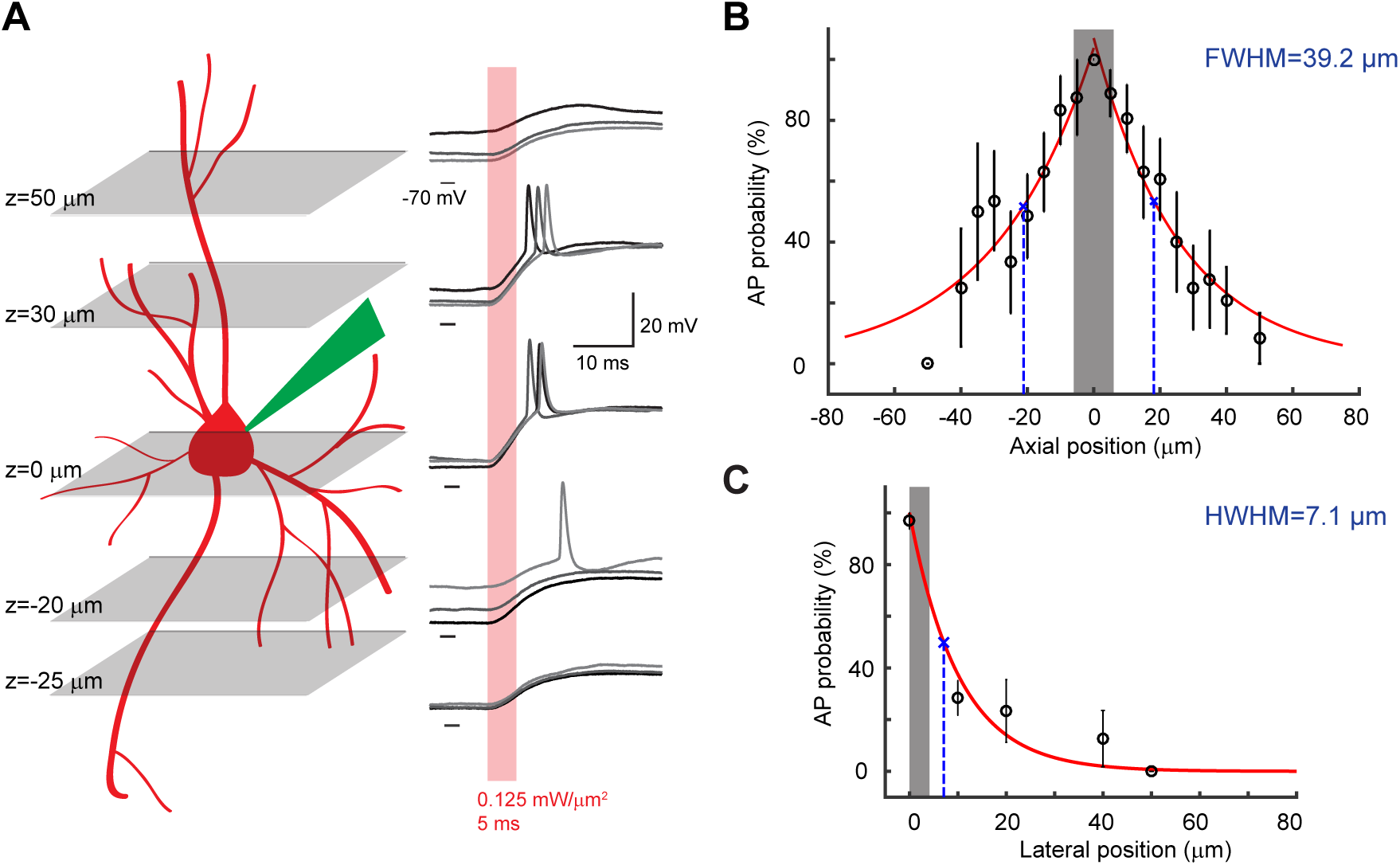
Spatial resolution of holographic activation *in vivo*. (A) Left: Schematic of mapping the axial resolution of holographic optogenetic activation in an opsin-positive cell by sequentially placing the excitation spot at different axial planes after manual defocusing. Right: Example whole-cell recordings of a ReaChR-expressing cells in response to 5-ms patterned illumination of 0.125 mW/μm2 at the corresponding axial positions. (B) Axial resolution of AP generation in response to photostimulation through a circular holo-graphic spot of 12-μm-diameter at the threshold excitation power in neurons (n=12) expressing ReaChR at L2/3 mouse visual cortex. Note the larger variability in AP generation when the excitation pattern is delivered deeper away from the focal plane. Shaded-grey area indicates the assuming axial span of target soma. (C) Lateral resolution of AP generation in ReaChR-positive cells (n=8) using the same photostimulation condition after placing the holographic spot laterally away from the target soma. Circles and error bars indicate mean±s.e.m. Grey-shaded area indicate the assuming lateral span of target soma.

### Concurrent holographic activation and optical read-out of multiple cells

To test whether holographic stimulation enables multi-cell activation in a neuronal network, we conducted simultaneous holographic activation of a neuronal subset while concurrently imaging the group calcium dynamics in mouse V1 *in vivo*. To this end, by injecting the opsin viral vectors of ReaChR-dTomato at L2/3 of V1 in the transgenic mice GP4.3 (32, 42), we co-expressed ReaChR and the calcium indicator GCaMP6s in a subset of cortical cells.

We confirmed the sensitivity of GCaMP6s in reporting single to tens of APs (32), by correlating the fluorescent signals with the spontaneous spiking activity measured using 2P-guided cell-attached recordings during imaging (Fig. S7A). Because GCaMP6s was expressed in almost all cortical neurons in transgenic mice, to identify double-positive cells of calcium sensor and opsin we only registered the spatial distribution of cells expressing ReaChR-dTomato in the red channel (Fig. 4A). In a field of view (FOV) of 300×300 µm^2^ with 66.4±4.6 identified neurons (19 FOV in 6 mice; 1 FOV was stimulated with 2 sets of spots), we selectively stimulated 7-9 cells through 12-µm holographic spots that were placed over target somata. The average distance between spot and the excitation field center was 44.6±1.2 µm (20 spot sets; Fig. S8). By stimulating with 10 light-pulses of 5 or 10 ms at 11.84 Hz (excitation power density between 0.05-0.3 mW/µm^2^, corresponding 7-34 mW/cell), and imaging at 5.92 Hz frame rate with a scanning power of 45.5±1.5 mW (20 experiments), we observed that target cells displayed fluorescent changes of GCaMP6s while most neighboring non-target cells did not show calcium events upon photostimulation (Fig. 4A). We found that with the increasing excitation power density, individual target cells displayed more evident calcium responses (Fig. 4B). The excitation intensity modulated both the activation probability, the ratio of the number of activated targets to the total number of targets (p<0.0001, 1-way ANOVA) and the peak amplitudes of evoked calcium transients (p=0.018). Compared to the stimulation condition of low excitation intensity 0.05 mW/µm^2^, the activation probability was significantly promoted when the excitation intensity increased to 0.1 mW/µm^2^ (p<0.0001, multiple comparisons between stimulation conditions), but remained of the same order of magnitude ~80% when the excitation intensity rose further. Likewise, the peak amplitudes of evoked calcium transients did not further increase when the excitation power density was risen above 0.2 mW/µm^2^ (Fig. 4C). Increasing the excitation intensity above the threshold of all target neurons would neither enhance the peak calcium response of individual target nor promote the activity probability of all targets, but rather deteriorate the axial confinement of light-patterns and could lead to excessive sample heating.

**Figure 4.**
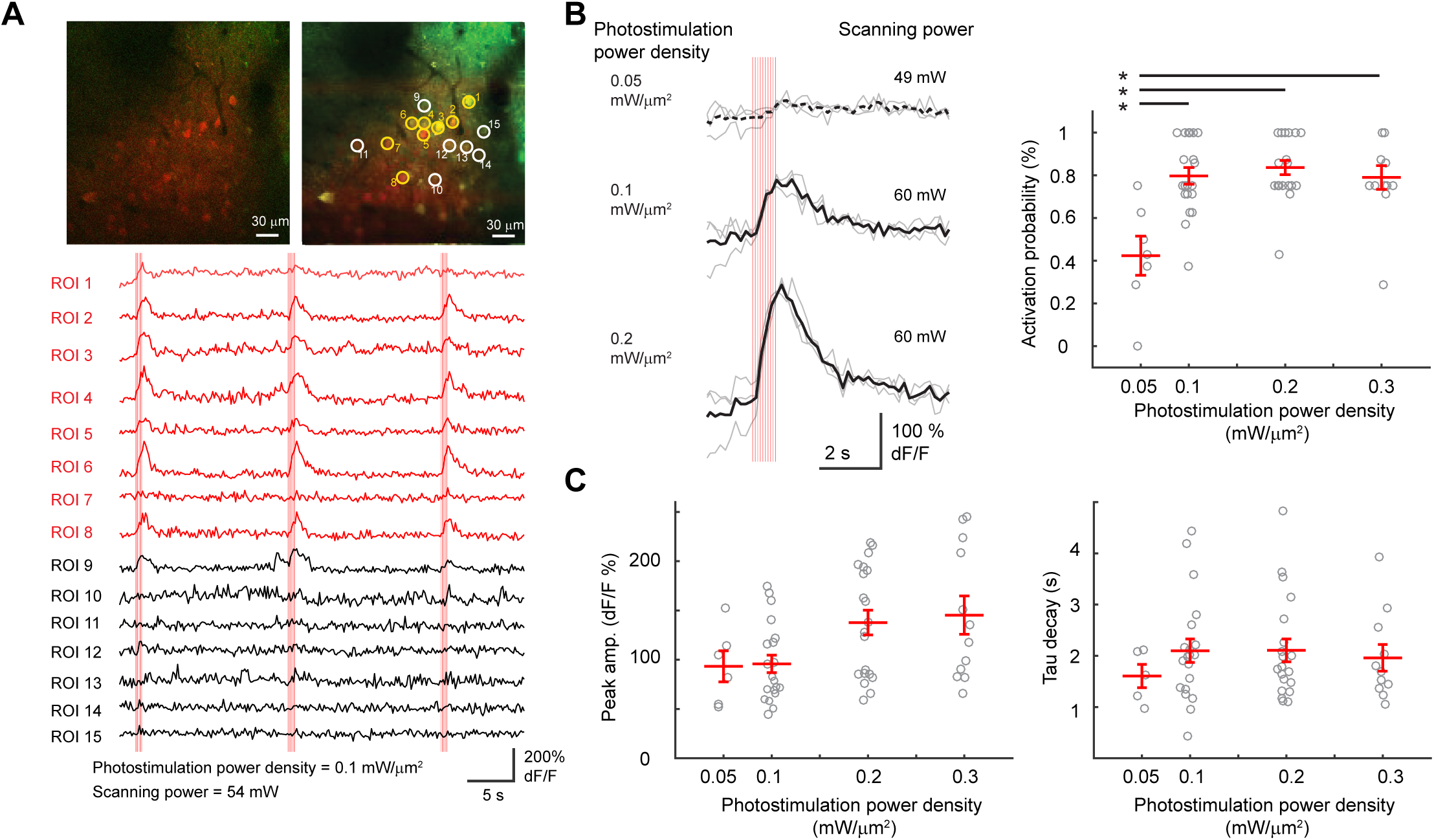
Concurrent holographic activation of multiple cells and optical readout of population activity. (A) Upper left panel displays a high-resolution 2P image of neurons co-expressing ReaChR-dTomato and GCaMP6s at the L2/3 of anesthetized mouse visual cortex. Upper right panel shows a maximum projection intensity profile from a low-resolution 2P image during all-optical experiment taken from the same FOV. 8 cells (right panel, yellow circles) were selected for simultaneous photostimulation through 8 holographic spots. While circles indicated out-of-target cells nearby. Bottom: Example calcium signal from the 8 target cells described above (red traces) and nearby neurons (black traces) in response to three epochs of photostimulation (vertical red bars). (B) Left panel showed example calcium signal from one cell, among 8 target cells, in response to photostimulation at different excitation intensities. Black traces represent the average from 3 repetitions (grey traces). Dashed line denoted insignificant photoactivation. Illuminations were indicated as vertical red bars. Right panel showed activation probability (mean±s.e.m.; 20 stimulation experiments) in relation to the excitation intensity. (C) Peak amplitude (left) and decay time constant (right) as a function of excitation power density.

By co-expressing ReaChR and GCaMP6s, we showed that TF-CGH enabled simultaneous multi-cell activation *in vivo*. ReaChR, an opsin of slow channel closing, however, is prone to cross-talk activation while scanning at high imaging power (Fig. S7B). In all-optical experiments, we therefore used a scanning imaging power of 30-60 mW to ensure clear visualization of calcium transients while inducing spurious AP firing < 3 Hz in ReaChR-positive cells. A better signal-to-noise ratio of evoked calcium responses may be attained by using a fast opsin, such as ChrimsonR, whose cross-talk activation is less severe upon high scanning power (Fig. S7C). Of note, 2P scanning may still lead to subthreshold activation in opsin-positive cells, thus rendering increased network excitability.

## Discussion

We used TF-CGH to demonstrate optogenetic activation at L2/3 of anesthetized mouse V1 with cellular resolution for three opsins of different channel kinetics, ReaChR, CoChR, or ChrimsonR. The enhanced peak power of laser-pulses from a fiber amplifier of low repetition-rate allowed suprathreshold photoactivation *in vivo* with moderate (in average ≤ 0.2 mW/µm^2^ or 23 mW/cell for the three opsins) excitation power density (33, 36, 43). The holographic pattern covers the entire soma of target neuron, thus enabling efficient current integration independently of opsins’ channel kinetics. Comparable with results of holographic photoactivation *in vitro* (33, 36, 43), we observed that higher illumination intensity led to shortened AP latency and reduced jitter, which was resulted from the promoted channel opening rate (44).

We found the photostimulation conditions which enabled AP generation with < 9 ms temporal resolution and < 1 ms temporal precision for these three opsins. The temporal properties of AP generation were preserved at high stimulation frequencies until 40 Hz for faster opsins of CoChR and ChrimsonR. For the slow opsin ReaChR, the combinatorial effect of delayed channel closing and inactivating lead to decreases in AP probability and spike’s temporal resolution and precision upon photostimulation > 20 Hz. Using the fast opsin ChrimsonR, we further demonstrated that the spontaneous activity can be precisely recalled in the target cell by designing a train of irregular light-pulses. Hence, holographic illumination can be applied for precisely reproducing a burst of spikes, synchronous firing, or mimicking spontaneous or evoked activity for closed-loop activity control in a neuronal ensemble of specific spatial organization (45, 46). Such neuronal spatiotemporal coding is of broad biological significance from synaptic mechanism (47, 48), plasticity (49, 50), sensory processing (51–53) to behavior (54, 55).

We showed that holographic illumination effectively activated opsin-positive neurons, here ChrimsonR, not only by viral infection but also via a novel transgenic mouse line for layer-specific labelling. The transgenic expression of opsin or activity reporter is useful for conducting all-optical control of multiple neurons, because it circumvents the issue of discrepant time windows for expressing two viral constructs. We performed *in vivo* all-optical experiments in double-positive cells by viral infection of ReaChR into V1 of the transgenic mice expressing GCaMP6s. Alternative combinations are viral expression of calcium indicator in the transgenic opsin line or double transgenic line.

In addition, the cytosol fluorophore expression initiated by the P2A sequence of opsin construct largely enhanced the visualization of positive cells (18, 30), thus providing clear reference for placing the holographic spots over target somata.

Parallel optogenetic activation using TF-CGH could be extended in several directions. First, by expressing soma-targeted opsins, un-biased neuronal connections could be identified by photostimulating one or many presynaptic cells while recording the postsynaptic responses with a glass microelectrode (33, 30) or using calcium imaging (56). Second, the functional reference of predetermined connections could be identified by correlating the performance of specific perception, such as the tuning of visual responses (57), or behavior (23). Finally, our current optical system could integrate a second SLM for generating axially-confined stimulation patterns in 3 dimensions (3D) (58, 59), which would be suitable for investigating the micro-circuitry between cortical laminae or brain regions. By further incorporating the 3D imaging methods (60–63), all-optical investigation of the functional wiring in a neuronal ensemble spanning a brain volume, for example a cortical column, may be realized.

## Materials and Methods

### Animals

All animal experiments were performed in accordance with the Directive 2010/63/EU of the European Parliament and of the Council of 22 September 2010. The protocols were approved by the Paris Descartes Ethics Committee for Animal Research with the registered number CEEA34.EV.118.12. Adult female or male C57BL/6J mice (Janvier Labs) were anesthetized with intraperitoneal injection of a ketamine-xylazine mixture (0.1 mg ketamine and 0.01 mg xylazine/g body weight) during stereotaxic injection and with isoflurane (2% for induction and 0.5-1 % for experiment) during photostimulation experiments. Adult mice of both sexes from the transgenic line GP4.3 (The Jackson Laboratory), which expressed the calcium indicator GCaMP6s (42), were used in all-optical experiment. Cortical neurons of 4-week-old mice were infected with viral vectors of opsins using stereotaxic injection. Holographic stimulation experiments were performed 5-12 weeks after injection.

### Virus injection and surgical procedures

For expressing ReaChR, CoChR or ChrimsonR, the following viral constructs were used respectively: AAV2/1-EF1α-ReaChR-tdTomato, AAV2/8-hSynapsin-CoChR-GFP, AAV2/8-hSynapsin-CoChR-mCardinal, and AAV2/7m8-CAG-ChrimsonR-tdTomato. Through a craniotomy over the right primary visual cortex (V1; 3.5 mm caudal from the bregma, 2.5 mm lateral from the midline), 1.5-2 µL viral vectors were delivered via a cannula in L2/3 (250 µm deep) at a speed of 80-100 nL/min. For performing acute photostimulation experiments *in vivo*, a circular craniotomy of 2 mm diameter was made over V1 and the dura mater was removed. Agarose of 0.5-2% and a cover glass were applied on top of the craniotomy to dampen tissue movement.

### Two-photon-guided electrophysiology *in vivo*

Neurons in L2/3 V1 of anesthetized mice were targeted with patch pipettes under a custom-built two-photon microscope equipped with a Ti:Sapphire laser (Chameleon Vision II, Coherent; pulse width 140 fs, repetition rate 80 MHz, average power at peak 3 W, tunable wavelength 680-1080 nm), and a 40X water-immersion objective (0.8 NA, CFI NIR Apo, Nikon). Details on the imaging setup are given in Fig. S1A. The fluorophore labelling of GFP, dTomato, or td-Tomato in cells expressing either of the 3 opsins or GCaMP6s was visualized by excitation at 920 nm and the emitted fluorescence was collected through red (617/70 nm) and green (510/80 nm) filters. Imaging data were acquired using ScanImage3 software (http://scanimage.org).

Whole-cell or cell-attached recordings were obtained by using microelectrodes fabricated from borosilicate glass (5-8 M Ω resistance) and filled with solution containing the following (in mM): 135 potassium gluconate, 10 HEPES, 10 sodium phosphocreatine, 4 KCl, 4 Mg-ATP, 0.3 Na3GTP, and 25-50 Alexa Flour 488 or 594 for pipette visualization. The craniotomy was covered with the extracellular solution containing the following (in mM): 145 NaCl, 5.4 KCl, 10 HEPES, 1 MgCl_2_, 1.8 CaCl_2_. Whole-cell membrane potential recorded in the current-clamp mode were corrected for liquid junction potential (11.7±0.3 mV, 16 measurements, mean±s.e.m.). Voltage recordings were acquired by using a MultiClamp 700B amplifier and a Digidata 1550A digitizer, which were controlled by a pCLAMP10 software (Molecular Devices). Electrophysiology data were filtered at 6 kHz and digitized at 20 kHz.

### Holographic stimulation *in vivo*

Holographic stimulation was implemented together with the imaging system mentioned above (Fig. S1A). Computer-generated holography was utilized for patterning light beams from an amplified fiber laser (Satsuma HP, Amplitude Systemes; pulse width 250 fs, tunable repetition rate 500-2000 kHz, gated from single shot up to 2000 kHz with an external modulator, maximum pulse energy 20 μJ, maximum average power 10 W, wavelength λ=1030 nm) operated at 500 kHz, via phase modulation through a liquid-crystal on silica spatial light modulator (LCOS-SLM X10468-07, Hamamatsu Photonics). The SLM was controlled by a custom-designed software, Wave-front Designer (21). Temporal focusing of the phase-modulated light pulses was performed through a reflective dispersion grating of 600 l/mm and a lens collimating the dispersed spectral frequencies at the back aperture of the 40× objective. The optical resolution obtained for a temporally-focused holographic spot of 12-μm diameter is characterized in Fig. S1B, and it was around 10 μm FWHM.

The experimental photocurrent-vs-illumination intensity for ReaChR published in Chaigneau et al. (36) can be fitted by an exponential function given by 
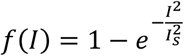
, where *I* is the illumination intensity and *I_s_* = 0.03 mW/μm^2^, corresponding to the intensity at 60% of photocurrent saturation. Hence, the axial photocurrent distribution will be given by

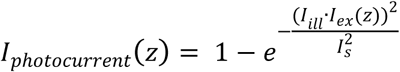

where *I_ill_* is the in-focus illumination intensity, and *I_ex_(z)* is the axial illumination profile. In our case, *I_ill_* has been estimated by calculating the average intensity used in the axial resolution experiments and assuming a scattering length at 1030 nm estimated as 175 μm, derived from simulations based on experimental data (26). The overall expected photoactivation axial distribution is then given by the convolution between the axial photocurrent distribution and the shape of the cell (Fig. S6A).

In anesthetized mouse V1, a circular holographic spot of 12-µm diameter was placed according to a high-resolution 2P reference image (512×512 pixels) including the target soma of an opsin-expressing cell. 2P photostimulation was performed via the holographic spot while the spiking activity was recorded through a patch pipette.

In all-optical experiments, target somata were mainly determined based on the ReaChR-dTomato expression. Through multiple 12-µm-diameter holographic spots, the targets were simultaneously illuminated while concurrently the population calcium activity in the 300×300 µm^2^ FOV was monitored via the fluorescence changes of GCaMP6s, which were recorded by using the imaging laser at 920 nm with 128×128 pixel resolution at a frame rate of 5.92 Hz.

### Data analysis

Electrophysiology recordings of single-cell photoactivation were analyzed using custom-written scripts in MATLAB (MathWorks). The latency of light-induced AP was defined as the time span between the illumination onset and the AP peak. The AP jitter was calculated as the standard deviation of light-induced AP latencies in 3-6 repetitions.

Image analysis was performed using ImageJ in combination with MATLAB. For all-optical experiments, region-of-interests (ROIs) covering individual somata were manually selected based on the expression of both ReaChR-dTomato and GCaMP6s channels. Relative percentage changes of fluorescence were computed for calcium signal as ∆F/F=(F-F_0_/F_0_), where F0 represented the average raw fluorescent signal of 1.5 s before photostimulation onset. A cell was considered activated when the mean ∆F/F 1.5 s after the first illumination onset was significantly different from that 1.5 s beforehand (paired t-test). Peak amplitude was determined as the peak ∆F/F signal within 1.5 s after the first illumination onset relative to the mean ∆F/F 1 s before the first light-pulse. Decay time constant of calcium event was determined in activated cells by fitting an exponential decay function to the average light-induced calcium transients till 2 s after the peak ∆F/F.

Statistical tests were conducted in MATLAB. Data comparisons between neurons for opsins types or different photostimulation conditions (e.g. frequency of a train of light-pulses, or excitation power density) were performed using ANOVA and multiple comparisons of Tukey’s method. Data between conditions (e.g. calcium signal before and after photostimulation onset in the same cell) were compared using paired t-test.

## Acknowledgements

The authors thank Edward S. Boyden and Deniz Dalkara for providing the viral constructs of CoChR and ChrimsonR viruses, respectively.

IWC received funding from the European Union’s Horizon 2020 research and innovation program under the Marie Skłodowska-Curie grant agreement no. 747598. EP and VE acknowledge the ‘Agence Nationale de la Recherche’ ANR (grants ANR-14-CE13-0016, Holohub and ANR-15-CE19-0001-01, 3DHoloPAc), VE acknowledge the Human Frontiers Science Program (Grant RGP0015/2016), the National Institutes of Health (Grant NIH U01NS090501-03) and the Getty lab.

